# Barley genetic architecture conditionally impacts recruitment of rhizosphere microorganisms and crop performance

**DOI:** 10.1101/2024.11.21.624704

**Authors:** Jessica L. Williams, Erik Z. Killian, Anna Halpin-McCormick, Michael B. Kantar, Jamie D. Sherman, Patrick M. Ewing, Jed O. Eberly, Jennifer Lachowiec

## Abstract

Microorganisms recruited to the rhizosphere from the surrounding soil can benefit the fitness of their host. Variation in plant genetics is associated with variation in rhizosphere microbial community composition leading to changes in fitness and crop productivity. However, what impact the abiotic environment has on connections between microbes and host genetics, and whether those connections in turn impact crop performance in realistic agricultural scenarios remains unclear. We assessed agronomic performance and 16S and ITS amplicon-based rhizosphere bacterial and fungal community composition on a large diverse barley population grown in seven field trials across four locations and two years. Within adapted regions, we observed consistent rhizosphere compositions across diverse soils, whereas in an unadapted environment, distinct microbial communities were recruited, indicating environmental specificity in microbial assembly. A Genome-wide association study (GWAS) identified 864 associations of barley genetic markers with bacterial or fungal taxa abundance. A total of 108 microbe-associated quantitative trait loci (QTL) co-localized with agronomic traits, suggesting pleiotropy or genetic linkage. Associated taxa varied considerably across field trial environments, whereas the mapped host QTL were more consistent. Members of the nitrogen cycling bacterial phylum, Nitrospirota were the most extensively associated. This phylum had 25 marker associations across five of the field trial location-years in the GWAS. These included a locus on chromosome 2H that associated with Nitrospirota bacteria and grain protein in two location-years, and the *Leohumicola* fungal genus and grain protein in a third trial. These findings support the genetic manipulation of rhizosphere microbiomes to enhance crop adaptation, whether with consistent or environment-specific microbial taxa. Such breeding advances will support phenotyping and selection strategies to improve crop resilience and productivity across variable environments.

## Introduction

Plant microbiomes contribute critically to plant survival and productivity. Microbes benefit plant health by facilitating the acquisition of nitrogen, phosphorus, and other nutrients (Pii et al. 2015; Rosenblueth et al. 2018), stimulating hormonal pathways to promote plant growth (Babalola et al. 2021), protecting against pathogens (Mendes et al. 2011) and mitigating abiotic stressors such as drought (Castrillo et al. 2017; Mayak et al. 2004; Poudel et al. 2021; Zhang et al. 2023). While the development of bacterial and fungal inoculants to enhance plant performance is an active area of research (Alori et al. 2017), their large-scale production, distribution, application, and establishment pose hurdles to their efficacy (Kaminsky et al. 2019). A different tactic that has received attention more recently is breeding crop varieties for an enhanced ability to recruit beneficial microorganisms (Clouse and Wagner 2021; Contreras-Liza 2021; Gopal and Gupta 2016). To do this, a detailed understanding of the interactions between plants and their associated microbiomes and how varying environmental conditions alter these interactions is needed. Thus, large knowledge gaps remain to be filled before microbial communities can be predictably manipulated via host genetic selection to benefit crop performance.

A critical zone ripe for microbial manipulation is the rhizosphere—the region surrounding plant roots that is characterized by dynamic interactions among diverse microorganisms and is considered one of the most complex biological habitats on earth (Ahkami et al. 2023; Chepsergon and Moleleki 2023). While soil microbial communities are driven by local climates and soil factors, the rhizosphere harbors a unique microbiome that is assembled from the soil microbiome and influenced by plant traits (Bulgarelli et al. 2013). The genetic basis underpinning plant control of rhizosphere microbiome assembly remains poorly understood (He et al. 2024; Trivedi et al. 2020).

Only traits under the influence of plant genetics can be targeted by selection in plant breeding. Some rhizosphere microbiome variation is attributed to plant genotype, revealing heritability of these extended phenotypes (Peiffer et al. 2013; Walters et al. 2018), suggesting breeding may be possible. Recent research has begun to uncover some of the associated loci through genetic mapping of root and rhizosphere microbiome extended phenotypes in domesticated and wild monocots and dicots, including *Arabidopsis thaliana* L. (Bergelson et al. 2019), *Sorghum bicolor* L. (Deng et al., 2021), *Panicum virgatum* L. (Sutherland et al. 2022), *Solanum lycopersicum* L. (Oyserman et al. 2022), and *Setaria italica* L. (Wang et al., 2022). These studies have minimized variation in the abiotic and biotic environment by providing a uniform starting soil microbial community, growing in a controlled environment, or examining a single field location and time point. This may aid in detecting the effects of plant genetics but overlooks the effects of important gene by environment (GxE) interactions that limit transferability of identified loci to non-experimental contexts.

Recent studies incorporating the examination of GxE on plant rhizospheres (Meier et al., 2022; Edwards et al., 2023) reveal the need to assess context to exploit the microbiome. Microbial communities are sensitive to a plethora of environmental factors such as water and nutrient availability, so the success of manipulating the rhizosphere microbiome for agricultural systems through crop breeding depends on incorporating plant-microbe-environment interactions (Bano et al. 2021). For example, the heritability of the rhizosphere microbiome is dependent on variation in local microbiota, and variation in soil type and fertility (Meier et al. 2022; Robertson-Albertyn et al. 2017).

Another influential but often overlooked aspect of the rhizosphere is the co-habitation of bacteria and fungi. Most microbiome studies have focused only on bacteria, but fungi can be just as impactful in their interactions with plants(Adedayo and Babalola 2023). Considering both of these kingdoms together adds a critical layer of understanding (Bergelson et al., 2019). Our previous research indicated that bacterial and fungal rhizosphere communities varied distinctly between plant subpopulations and in response to environmental factors (Killian et al. 2026).

To understand GxE interactions shaping both bacterial and fungal members of the rhizosphere microbiome, we used a barley (*Hordeum vulgare* L.) population representative of five different breeding programs based in Idaho, Minnesota, Montana, North Dakota, and Washington: The Spring Two-Row Multi-Environment Trial panel, S2MET (Neyhart et al. 2019). Due to its history of selection, this population contains both lines that are locally adapted and lines that are more generally adapted across the growing regions (Ewing et al. 2024). We grew this population in field trials within and far outside the adapted range of barley to investigate connections between barley genetics, rhizosphere microbial community characteristics, and local adaptation. We explore the colocalization of microbiome associations with agronomic traits and discuss how the current research informs the long-term goal of breeding for plant-microbe interactions that improve crop adaptation.

## Materials and Methods

### Plant materials and genotyping

We explored the S2MET Barley Mapping Population to understand associations between plant genotype and microbiome, and implications for plant performance. The S2MET panel includes lines from breeding programs at USDA-ARS Aberdeen in ID, University of Minnesota, Montana State University, North Dakota State University, and Washington State University. It was previously genotyped by sequencing (Neyhart et al. 2019) and mapped to the MorexV1, IBSCv2 barley reference genome (Mascher et al. 2017), and genotyping data is publicly available (Neyhart and Smith 2019). After matching the genotyping data to the subset of S2MET lines that were included in the current experiments, markers with a minor allele frequency (MAF) less than 5% were removed from the dataset. This left 4,787 single nucleotide polymorphism (SNP) markers that were used in the genome-wide association study (GWAS).

### Experimental design

Fields trials were performed in 2021 and 2022 at the Post Research Farm in Bozeman, MT (southwest Montana-swMT), the Central Agricultural Research Center in Moccasin, MT (central Montana-cntrMT), Brookings, South Dakota (SD), and in 2021 in Kunia, Hawaii (HI) (**Table S1**). An augmented randomized block design (Federer 1961) was used for the field trials in which each of the 232 experimental lines appeared once in the trial at each location, and four barley check varieties ‘Odyssey’, ‘Hockett’, ‘Lavina’, and ‘Merit 57’ appeared at random positions in each of twelve blocks. Seed was treated for broad range pathogen protection prior to planting and weeds were managed with herbicide according to regional agronomic practices. All trials were rainfed. Climate conditions including in-season precipitation, and soil profiles including nitrogen levels are reported in Supplemental Table S1.

### Plant phenotyping

The barley population was assessed for various agronomic traits in the field trials. Heading and maturity dates, stages 59 and 89 on the Zadoks cereal growth scale (Zadoks et al. 1974), were noted when seed heads were visible for half of the plants in a plot and when half of the seed heads had lost green color respectively. Duration of the grain fill period was calculated by subtracting heading date from maturity date. Plant height was measured after maturity from the base of the plant to the top of the seed head, excluding awns. Also after maturity was complete, three 15 cm sections were hand cut from each plot and weighed for total biomass. Seed heads from these samples were counted to measure productive tiller number and were removed and threshed. The weight of this seed was divided by the total biomass to calculate harvest index. Full plots were harvested by combine after grain ripening, and seed was weighed for gross yield calculations. Grain protein and moisture content were measured using near-infrared spectroscopy (Infratec NOVA, FOSS, Hilleroed, Denmark). Kernel plumpness was measured as the percentage of kernels retained on a 6/64 × 3/4-inch sieve following 30 seconds of shaking on a Pfeuffer Sortimat (Pfeuffer GMBH, Kitzingen, Germany). Grain test weight was measured using a John Grain Analysis Computer 2500-UGMA (Corn Belt Testing Inc.; Minneapolis, MN, USA). In Hawaii accessions remained vegetative, thus only total biomass was measured.

### Rhizosphere sampling and 16S and ITS amplicon sequencing

Amplicon sequencing to determine rhizosphere microbial community composition was previously described (Killian et al., 2026). In brief, the rhizosphere was sampled at early stem elongation and six plants with intact roots were removed from each plot for a composite sample. For rhizosphere isolation, plant roots with attached soil samples were vortexed in a 0.9% NaCl solution and sonicated in a water bath for 1 min. The isolated soil solution pellet was resuspended in sterile deionized H_2_O and stored at −70°C. DNA was purified from rhizosphere extracts and bulk soil using a QIAGEN DNeasy PowerSoil Pro DNA Isolation Kit (QIAGEN Inc., Germantown, MD, USA). Purified DNA was submitted to Integrated Microbiome Resource (Dalhousie University, Halifax, NS) for library preparation and amplicon sequencing of the bacterial 16S V6-V8 and fungal ITS2 regions. Sequencing was performed on the Illumina MiSeq platform (Illumina, San Diego, CA, USA) using a 2 300 paired end kit.

### Microbial community analysis

Quality filtering, trimming, chimera removal, and assembly of reads into error-corrected amplicon sequence variants (ASVs) were performed in R using the Dada2 package (Bolyen et al. 2019; Callahan et al. 2017). First, reads were trimmed to remove low-quality sequences. Reads were trimmed by 5 bases from the beginning of each read, then each location-year was trimmed to 200 bases (Kyle and Klassen 2024; Pauvert et al. 2019). Only forward reads were used for downstream analysis due to low-quality reverse reads. In all location-years filtering and trimming were performed with the default settings for the Dada2 pipeline. Due to the low read quality of the swMT21 fungal samples, they were excluded from subsequent analysis. Demultiplexed reads from all location-years were merged as sequence tables before aligning with field and taxonomy data. Taxonomic assignment was performed using the SILVA v138.1 SSU database with a 99% identity threshold (Quast et al. 2012).

Statistical analyses and data visualization were performed using phyloseq (McMurdie and Holmes 2013), microeco (Liu et al. 2021), and ggplot2 (Wickham 2016) packages. Relative abundances of major taxa were visualized at the phylum level using a 1% average relative abundance threshold. To gain insight into differences between rhizosphere communities across different environments, we performed principal coordinate analysis using centered log-ratio (CLR) transformed Aitchison distances (Lozupone et al. 2007). A supervised random forest machine learning approach was used to identify indicator taxa associated with specific locations (Paes da Costa et al. 2024) and the top 20 taxa based on Mean Decrease Gini (MDG) were plotted.

### Phenotypic Data Preparation for Genetic Analysis

Relative abundances of bacterial and fungal taxa in the rhizosphere samples were considered microbial traits of the plants and were prepared for analysis as follows. Datasets from location-year field trials and either 16S or ITS sequencing results were prepared and analyzed separately. To account for data sparsity, using the phyloseq R package (McMurdie and Holmes 2013), ASVs were filtered to only retain sequences that appeared in at least 0.7% of barley plot rhizosphere samples (removing singletons) and had a relative abundance (count divided by within-sample count total) of at least 0.1% in at least one sample. ASV read counts were then agglomerated at the genus rank. Zero counts were imputed using the Geometric Bayesian multiplicative (GBM) method in the cmultRepl function of the zCompositions package (Palarea-Albaladejo and Martín-Fernández 2015). This replacement of zeros (Quinn et al. 2019) allowed the data to then undergo CLR transformation using the Compositions R package, to account for the compositional nature of the data (Van den Boogaart and Tolosana-Delgado 2008). Agglomeration, zeros imputation, and CLR transformation were repeated at each taxonomic rank up to phylum. Taxa were eliminated from further processing and analysis if they did not appear on the list of prevalent taxa for their location-year and taxonomic rank. These prevalent taxa were ones that had a count of at least one in at least 80% of samples when looking at agglomerated raw counts.

Variance was partitioned using the R package, lmerTest (Kuznetsova et al. 2017) by fitting the following linear model for each taxon abundance:

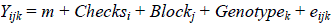

where Checks_i_ was a fixed effect representing the replicated check varieties, Block_j_ was a random effect, and Genotype_k_ was a random effect representing the experimental barley lines from the S2MET population. The percentage of the total variance due genotype was calculated to determine heritability (Holland et al. 2003). The percentage of total variance due to block was also determined and if this value was greater than 0, Best Linear Unbiased Predictions (BLUPs) were calculated for the trait to account for spatial variation within the fields by utilizing the augmented experimental design (Kehel et al. 2010). Coefficients of the Genotype_k_ factor corresponding to each barley line were the BLUPs used as input for GWAS analysis. If the Block effect was 0, CLR transformed values were used for genetic mapping of the trait without calculating BLUPs. The workflow for preparing microbial trait data for the GWAS is summarized in **Fig. S2.**

### Genome Wide Association Studies

Genetic mapping was carried out within each location-year and either agronomic, 16S, or ITS dataset in the program GAPIT using the FarmCPU model(Wang and Zhang 2021). Two principal components were used to account for population structure which appeared to provide appropriately fit models for traits when examining QQ plots. A Benjamini-Hochberg procedure was used to determine false discovery rate (FDR) adjusted p-values for marker-trait associations (Benjamini and Hochberg 1995). A significance threshold of FDR adjusted p < 0.2 was used to explore the results in order to avoid false negatives. For all microbial trait SNPs passing the significance threshold, Wilcoxon’s signed rank tests were also performed to compare the effects of the major and minor alleles (homozygous) on each agronomic trait. Significance for these was determined with a Bonferroni correction (P < 2.3033 ×10^-6^).

## Results

### Environmental variation shaped rhizosphere microbiomes and agronomic traits in barley’s adapted and unadapted ranges

We investigated the effect of location, within and outside the adapted barley range to uncover the interaction between host genetics and environmental variation in shaping rhizosphere bacterial and fungal communities. Field trials containing 237 S2MET barley genotypes for two years in three locations represented the adapted range of barley in the United States, including southwest Montana (swMT), central Montana (cntrMT), and Brookings, South Dakota (SD). We also conducted a trial for one year in Kunia, Hawaii (HI) which represents an unadapted region where barley is not grown. The field locations had distinct climates and soil physical and chemical properties (**Table S1**).

Within the adapted range, rhizosphere bacterial and fungal community composition was relatively similar, and contrasted with those of lines grown in the Hawaii environment (**Fig. 1a-f**). Actinobacteriota, Proteobacteria, and Bacterioidota were the dominant bacterial phyla associated with the barley rhizosphere and represented around 80% of the relative abundance at the adapted locations, which is consistent with other plant rhizosphere studies (Eberly et al. 2024; Ling et al. 2022; Zhang et al. 2024). Samples from Hawaii were also dominated by Actinobacteriota and Proteobacteria, but the relative abundance of Actinobacteria was lower (Fig. 1A). The dominant fungal phyla in the adapted range were Ascomycota and Mortierellomycota, similar to other barley rhizosphere studies (Liu et al. 2022; Ye et al. 2022), while Hawaii rhizosphere fungal communities had reduced representation of Mortierellomycota.

**Fig. 1.**
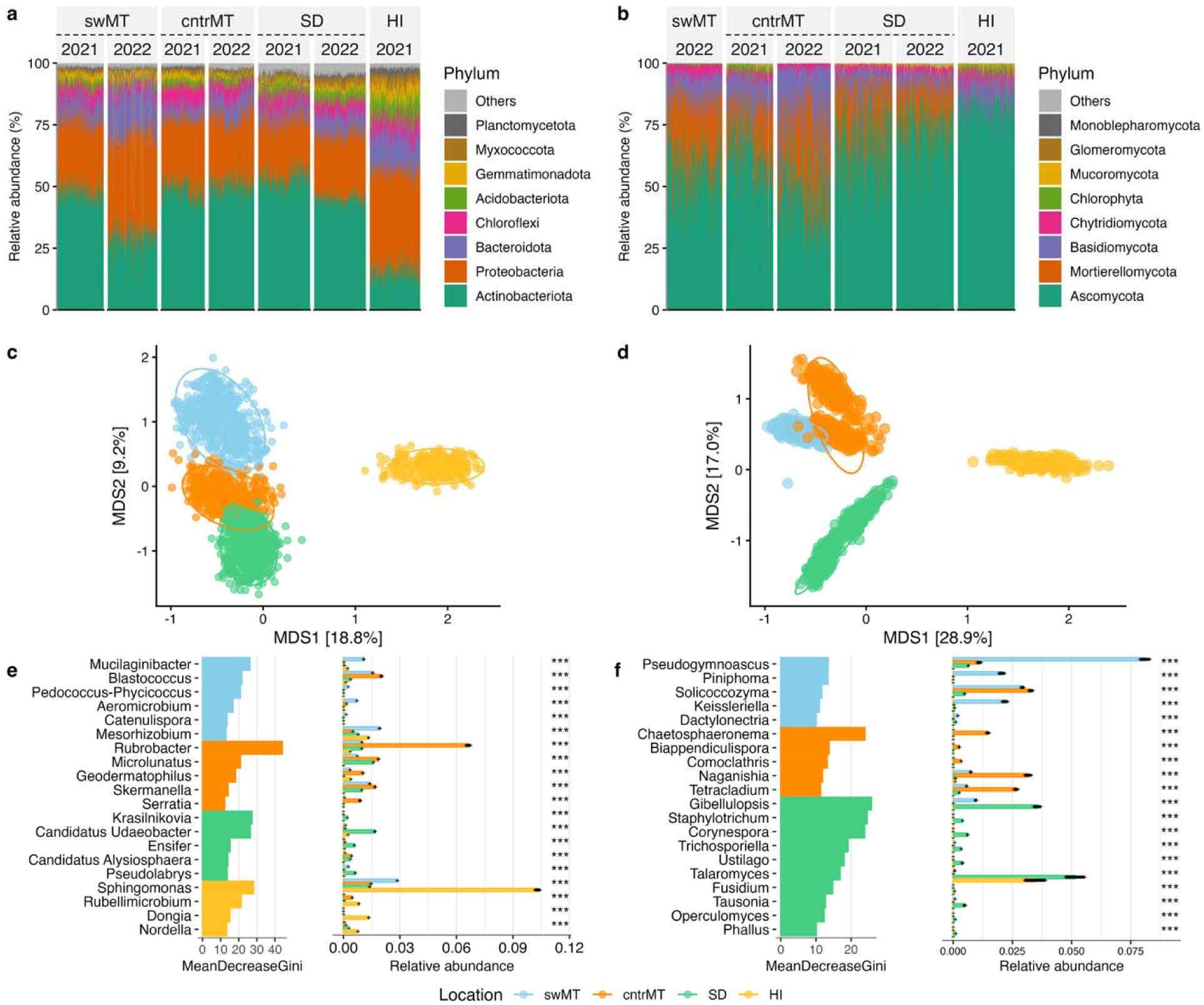
Location-specific variation in rhizosphere microbiomes across barley field trials. Relative abundance of bacterial (a) and fungal (b) phyla are shown for each location-year. Multidimensional Scaling analysis of centered log-ratio transformed abundance by genera in the rhizosphere bacterial (c) and fungal (d) communities among locations are shown. Differentially abundant bacterial (e) and fungal (f) genera predicted by a random forest model with Mean Decrease Gini (MDG) based on transformed abundances. swMT: southwest Montana shown in light blue, cntrMT: central Montana shown in orange, SD: Brookings, South Dakota shown in green, HI: Kunia, Hawaii shown in yellow.

Within the adapted locations, rhizospheres at individual locations had key differences in taxonomic compositions (**Fig. 1c, d**). Several bacterial and fungal genera were found to be differentially enriched in one location relative to the others (**Fig. 1e, f, Table S2**). For example, cntrMT had relatively high Rubrobacter and Chaetosphaeromena abundance relative to the other locations, whereas swMT showed higher abundances of Mucilaginibacter and Pseudogymnoascus while Sphingomonas was nearly 3-fold more abundant in HI compared to the other locations.

In addition to the differences in rhizosphere microbiome composition, we observed large differences in plant traits among the field trials, including phenology, shoot and grain morphology, and yield (**Fig. S1**). The photoperiod in Hawaii prevented flowering of any of the lines and thus agronomic assessment, consistent with the lack of barley adaptation to the location.

### Evidence for shared genetic determinants of microbial traits

We next investigated the contribution of barley genetic variation to rhizosphere microbiome assembly. We considered the bacteria and fungi sequenced from the rhizospheres as extended phenotypes (Whitham et al. 2003) of each barley line. We defined these extended phenotypes or ‘microbial traits’ as the centered log-ratio transformed abundances of the prevalent taxa (observed in at least 80% of lines) found in each location-year, calculated at each taxonomic rank from phylum to genus (**Fig. S2)**. A total of 769 microbial traits (324 bacterial, 445 fungal) and 12 agronomic traits across the seven location-years were explored. The heritability of the 2,410 total microbial trait by location-year combinations (1,365 bacterial, 1,045 fungal) ranged from 0.000 to 0.819 for bacterial traits and to 0.738 for fungal traits, where 14% and 13% had heritability of 0.3 or greater and 46% and 47% had 0 heritability for bacteria and fungi respectively (**Table S3**). The microbial and agronomic traits were mapped in a Genome Wide Association Study (GWAS). There were instances where a microbial trait had very low heritability but still had significant marker associations in the GWAS. This could arise because heritability was calculated using unreplicated barley lines in our augmented experimental design, while the GWAS model considers replicated alleles throughout the population (Edwards et al., 2022).

The patterns of quantitative trait loci (QTL) across the barley genome and among locations were consistent with pleiotropy or linkage, and GxE underpinning the genetic architecture of microbial traits. There were 864 associations between specific SNPs and microbial traits (562 bacterial, 302 fungal, **Table S4**). Of the bacterial taxa included in the GWAS, 213 (15.6%) displayed GWAS hits over our significance threshold while this number was 129 (12.3%) for fungal taxa. We identified 202 genetic hotspots where consecutive microbial trait associations were not more than 1Mbp apart (Karki et al. 2024) (**Fig. 2a, Table S4**), representing potential pleiotropy or tight genetic linkage among microbial traits. Although most taxa passed the prevalence threshold for mapping in more than one location-year (75% for bacteria, 56% for fungi) (**Fig. 2b, c**), it was much less common for a taxon to have GWAS hits in more than one location-year (29% for bacteria, 15% for fungi), and no taxa had a hit in every field trial (**Fig. 2d, e)**. Genetic hotspots, on the other hand, were more repeatable across location-years, particularly for bacterial GWAS hotspots (54% of which appeared in more than one location-year), though less so for fungal (25%) (**Fig. 2f, g)**. Prevalent taxa and GWAS results were not more similar at the same location over different years than at different locations. (**Fig. S3**).

**Fig. 2.**
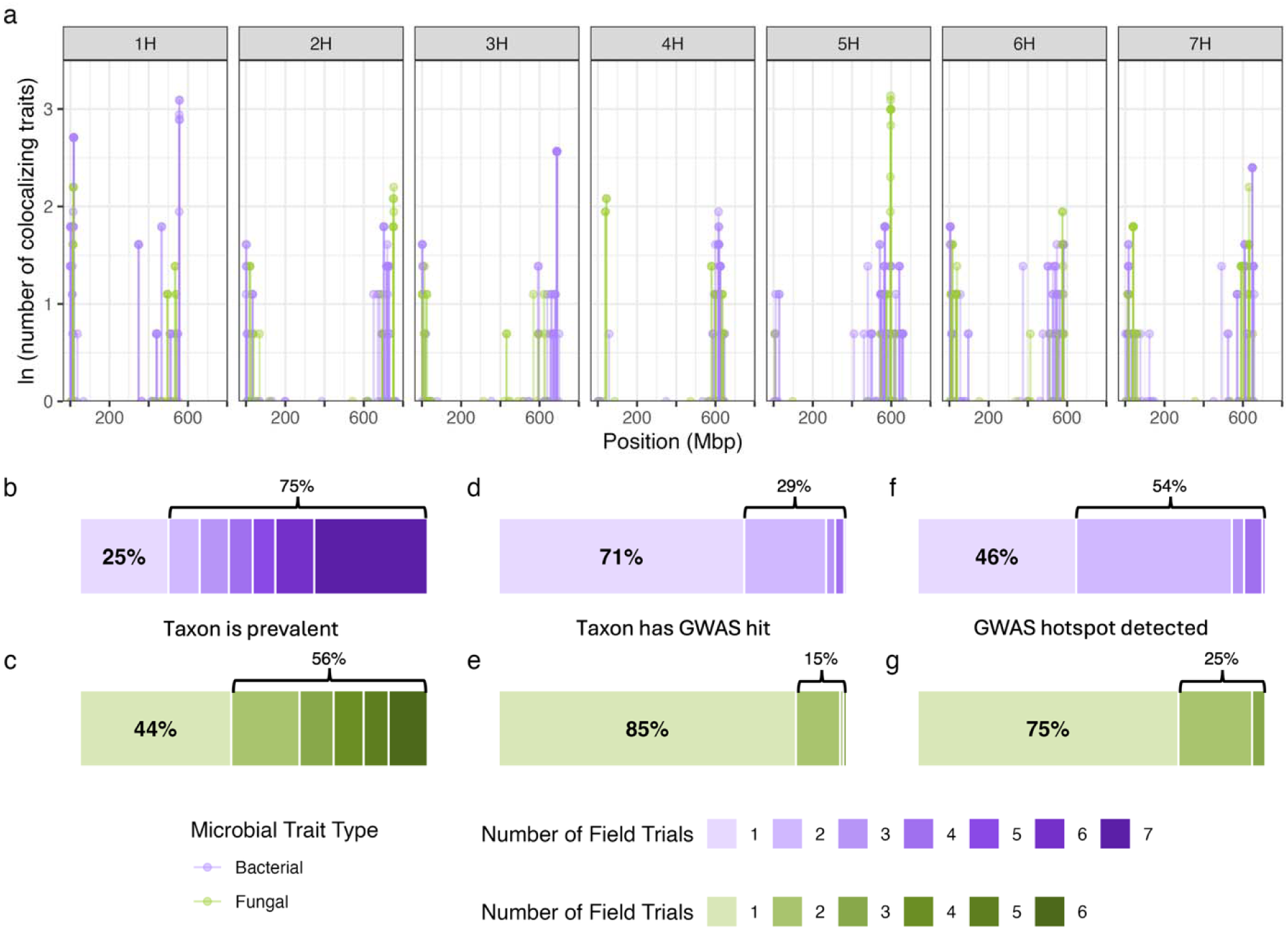
GWAS reveals genetic architecture of microbial and agronomic traits across field trials. (a) The natural log of the number of microbial trait associations (from any location-year) within 1Mb of the genomic position is shown. Bacterial traits colocalizing with bacterial traits are in purple and fungal with fungal in green. Of the bacterial (b) and fungal (c) taxa that passed the prevalence threshold for inclusion in the GWAS as microbial traits, the percentage that passed this threshold in 1 up to all 7 (bacteria) or 6 (fungi) location-year field trials is shown. Of the bacterial (d) and fungal (e) taxa that had any GWAS hits, the percentage that had hits in 1 up to 7 or 6 field trials is shown. Of the bacterial (f) and fungal (g) associated genetic hotspots, the percentage that appeared in 1 up to 7 or 6 field trials is shown.

While environmental effects dominated the taxonomy of associated bacteria and fungi, there were reproducible genetic signals with potential relevance for crop performance. There were 17 instances in which microbial traits from the same phylum mapped to the same genetic hotspot in multiple location-years, 11 of which at the class rank, and 2 of which matched at the order rank There were no instances of microbial traits from the same family or the same genus mapping to the same hotspot in more than one location-year. Never did the same exact taxon map to the same exact SNP in more than one field trial, but there were examples of different taxa mapping to the same loci, and the same taxon mapping to different loci across trials. Examination of repeated genetic hotspots revealed diverse microbial taxa with similar plant beneficial functions mapping to the same regions in different location-years (**Table 1**).

**Table 1.**
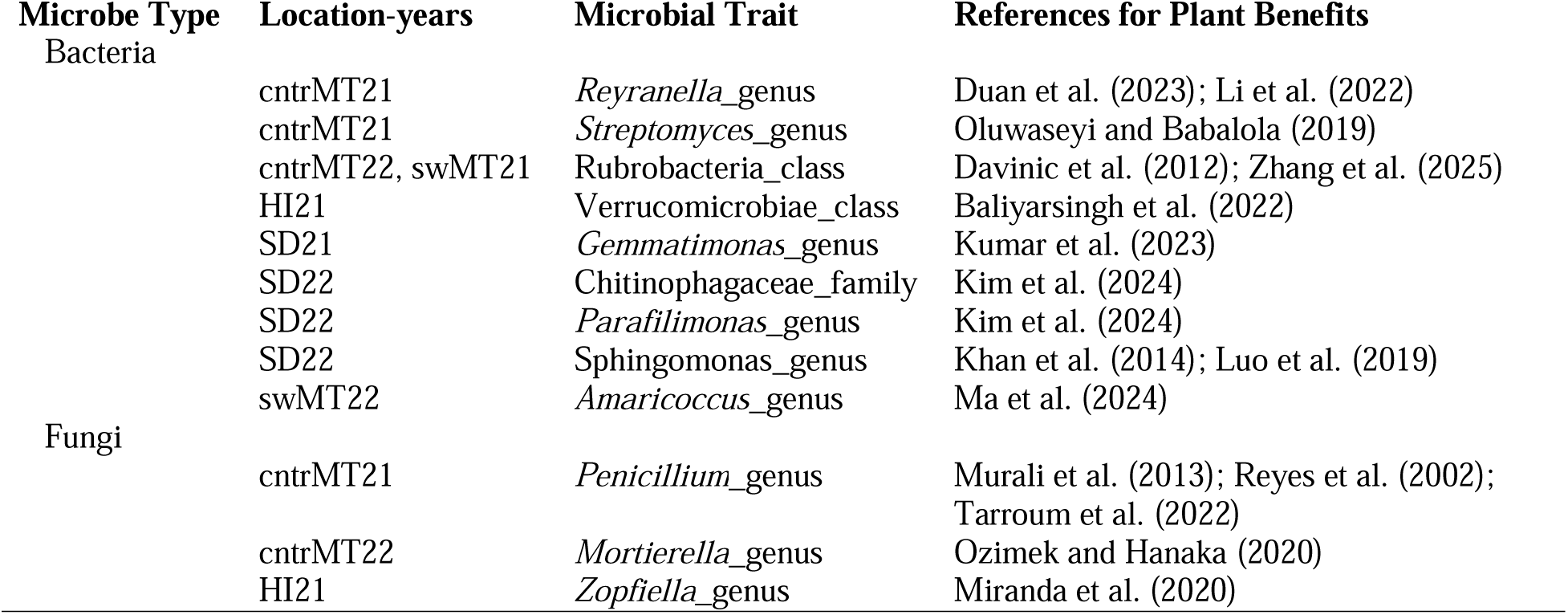
Summary of one of the genomic regions associated with multiple plant beneficial microbes in the GWAS results. Each of the seven location-years was represented here. Most of these microbial traits have been reported to have possible plant beneficial functions including plant growth promotion and nutrient cycling, abiotic stress tolerance, and biocontrol. All were associated with at least one SNP within an 8.6Mb region on chromosome 1H (8,973,024 – 17,610,771 bp) with some traits mapping to the same SNP or 1Mb hotspot within this region. Exact positions are given in Supplemental Table S3.

One taxonomic group stood out as being most sensitive to barley genetics based on its number of GWAS associations and representation in the GWAS results of multiple field trials. While the average number of GWAS hits for bacterial traits was 3.7, the Nitrospirota phylum had the most at 25 and each rank within that phylum also had abundant GWAS associations (**Fig. 3a**). This bacterial phylum in our study contained a single class (Nitrospiria, 15 hits), order (Nitrospirales, 22 hits), family (Nitrospiraceae, 21 hits), and genus (*Nitrospira*, 11 hits). The Nitrospirales order had GWAS hits in five of seven location-years, while no bacterial taxon outside of the Nitrospirota phylum had hits in more than three location-years. In swMT22, seven loci were associated with Nitrospirota phylum members, and when considering the effect sizes of these associated loci on the transformed abundance of *Nitrospira* (**Fig, 3b-g**), we observed an additive effect of the positive alleles (**Fig. 3h**). Certain Nitrospirota bacteria oxidize ammonium to nitrate, which is more readily available to plants (Koch et al. 2015).

**Fig. 3.**
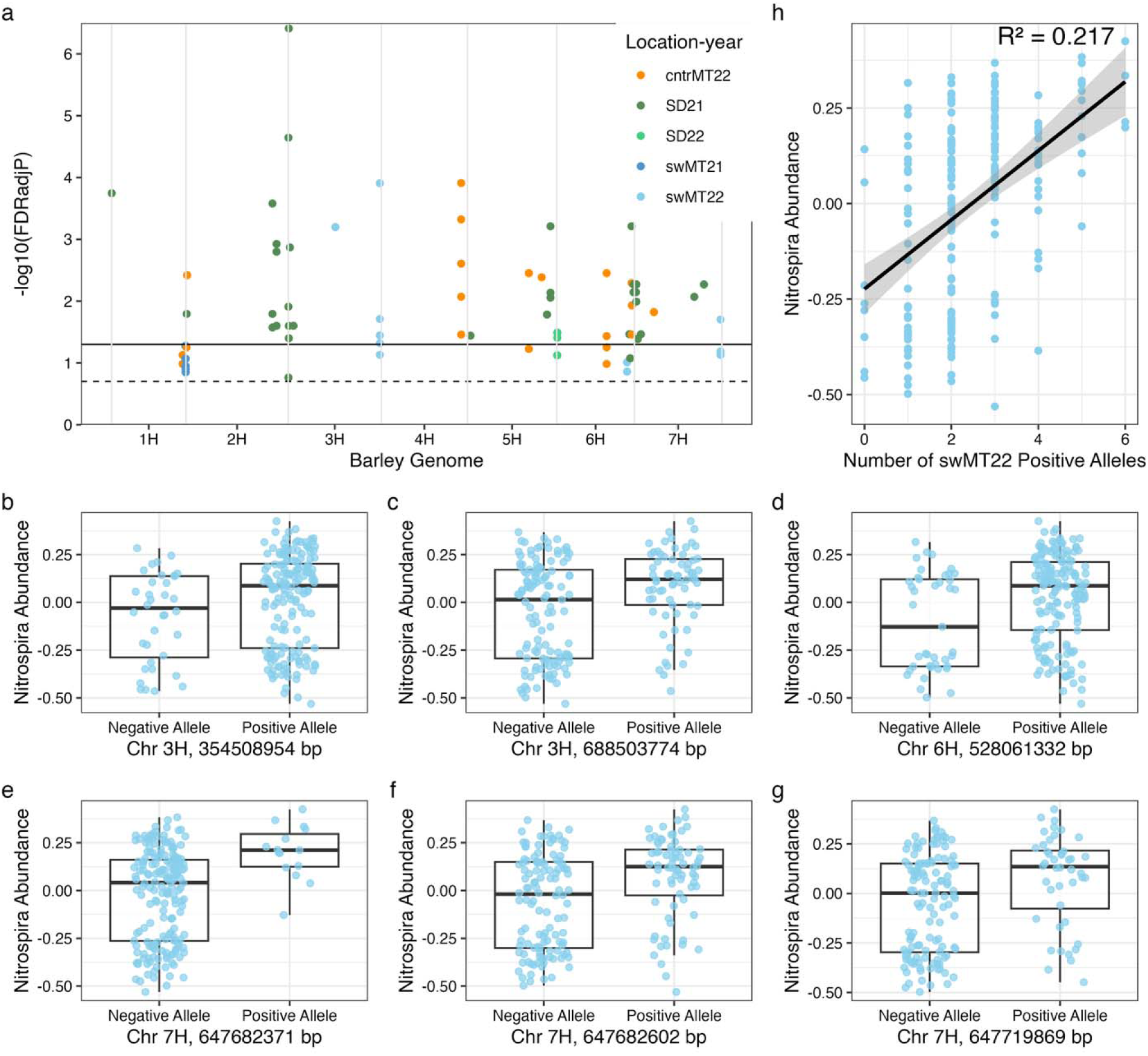
Genetic architecture associating with the Nitrospirota phylum. (a). All significant marker-trait associations for bacterial traits within the Nitrospirota phylum (within which there was only one class, order, family, and genus in this dataset). The dashed line indicates the exploratory FDR adjusted p < 0.2 threshold and the solid line is at FDR p < 0.05. Colors indicate the location-year. (b – g) Effects plots comparing transformed abundances of the *Nitrospira* genus in swMT22 between allelic groups at each SNP associated with bacterial traits from the Nitrospirota phylum in that location-year. (h) Additive effect on the transformed abundance of the *Nitrospira* genus in swMT22 of the number of positive alleles associated with the Nitrospirota phylum or its members in that location-year.

For fungal traits, the average number of hits was 2.7 and the *Rhizophlyctis* genus had the most hits with 12, which came from only two location-years. The most widespread fungal taxon was the Eurotiales order which had a total of seven GWAS hits in four out of six location-years. The genetic architecture affecting microbial traits was highly quantitative (many loci) and environmentally sensitive (different loci in different trials).

### Substantial overlap of genetic architecture contributing to microbial and agronomic traits

Of particular interest were microbial trait loci that also associated with agronomic traits, as this colocalization would be consistent with a role of the microbial taxon influencing plant growth and facilitating adaptation (Bulgarelli et al. 2015). In addition to scanning GWAS results for microbial and agronomic trait associations within 1Mbp of each other (Karki et al. 2024), we also used Wilcoxon’s signed-rank tests to examine whether microbial trait QTL alleles were associated with agronomic traits. Together, these approaches uncovered 1,273 significant marker associations with agronomic trait by location-year combinations in the adapted environments after multiple testing correction (Escudero-Martinez et al. 2022).

Bacterial and fungal taxa that were key to distinguishing one location environment from the others in our broad microbiome characterization (**Fig. 1e, f**) were also represented in our GWAS results and appeared related to location-specific crop performance in some instances. For example, the *Paenarthrobacter* bacterial genus, which was enriched for the SD environment, also mapped to a SNP on chromosome 6H in the SD21 trial. This locus was further implicated for grain protein content, grain plumpness, heading date, and grain fill duration in SD21 but no other location-year.

There were 108 instances in which an agronomic trait colocalized to the same 1Mb hotspot as a bacterial or fungal trait from the same location-year field trial (**Fig. 4a**). Of these hotspots, 23 contained such a match for two trials, and another three contained a match for three trials. One of these hotspots on chromosome 2H was of particular interest for its microbial trait associations in three location-years with grain protein content, a key quality trait in malt barley (**Fig. 4b**). It was not more common for these matching trials to be from two years at the same location (7 out of the 26 total). Thus, the overlap of microbial and agronomic genetic associations was highly influenced by environmental factors stemming from both location and trial year.

**Fig. 4.**
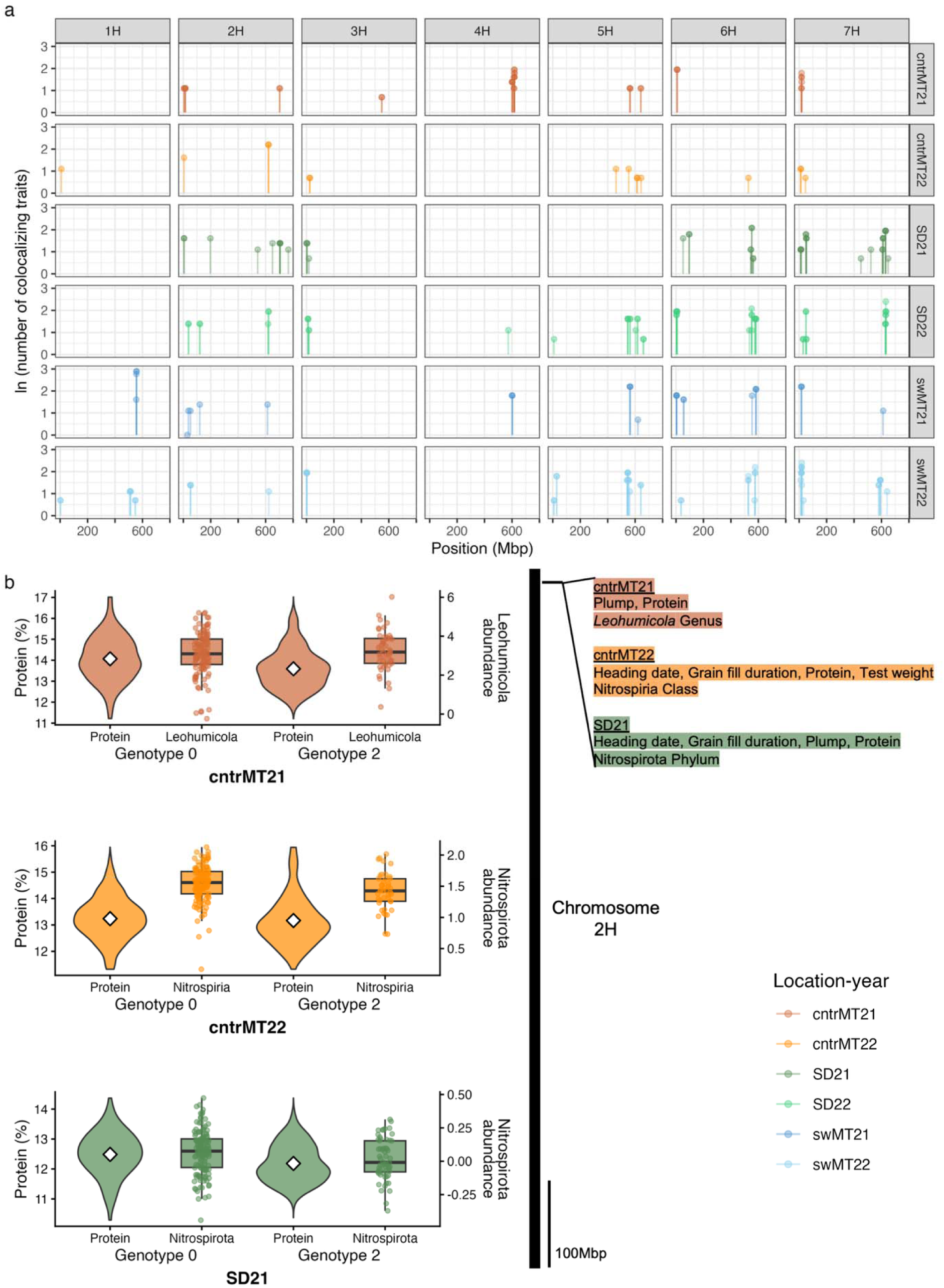
GWAS reveals overlapping genetic architecture of microbial and agronomic traits across field trials. (a) Lollipops indicate the natural log of the number of colocalizing trait associations (from GWAS or Wilcoxon’s test) within 1Mbp of the genomic position with the requirement that at least one microbial trait and one agronomic trait from the same location-year be present. (b) Microbial and agronomic traits from three location-years colocalizing to 5,023,518 bp on chromosome 2H. Violin plots compare grain protein distributions between allelic groups in the different field trials and boxplots represent the corresponding microbial abundance. Genotype 0 refers to the homozygous reference allele and Genotype 2 refers to the homozygous alterative allele.

Broader genetic regions did contain clusters of agronomic-microbial hotspots from more field trials. A ∼34Mb region on the short arm of chromosome 7H contained within-trial agronomic-microbial trait colocalizations from each of the six location-years within the adapted range. Leveraging such connections between microbial traits and agronomic traits may advance genetic and agronomic gain within target environments (Wang et al. 2022).

## Discussion

We have expanded our understanding of plant genetic relationships with the rhizosphere microbiome, evaluated the association of those relationships with agronomic traits targeted by breeding, and explored the role that environment plays in those interactions. Previous studies have demonstrated that plant recruitment of rhizosphere microorganisms varies with environmental factors (Edwards et al. 2023; Khasanova et al. 2023). We provide insight into the genetic variation that enables microbiome recruitment across and within environments. Consistent with local soil environmental effects (Crowther et al. 2019; Delgado-Baquerizo et al. 2018), barley recruited distinct rhizosphere communities across and outside of its adapted range that nonetheless were locally consistent across years (**Fig. 1c, d**). Rhizosphere compositions were mappable (**Fig. 2a**) and colocalized with plant traits (**Fig. 4**) suggesting adaptation to local conditions. Together, these findings are consistent with the selection of microbial communities for specific environments to maximize plant productivity (Schnitzer et al. 2011).

Since rhizosphere microbiomes assembled from the surrounding soil impact plants, and plants influence this assembly (Bergelson et al. 2019; Deng et al. 2021; Walters et al. 2018), recruitment of beneficial microbiomes through plant breeding is a logical next step. The sensitivity of the microbial community structure to environmental factors and interactions within the community makes genetic dissection challenging (Bergelson et al. 2019), and suggests that recruitment of specific taxa might not be consistent across environments. Nonetheless, we identified microbial associated genetic loci (**Fig. 2a**) in seven field environments (**Table S1**). Most taxa only mapped in a single field trial (**Fig. 2d, e**), indicating GxE that could contribute to local adaptation (Edwards et al. 2023; Meier et al. 2022; Petipas et al. 2021; Revillini et al. 2016). However, many GWAS hotspots had associations in multiple field trials (**Fig. 2f, g)**, particularly for bacteria, though rarely with the same microbe, potentially reflecting high levels of functional redundancy among soil-born bacterial taxa (Louca et al. 2018) (**Table 1**). Loci that mapped repeatedly over multiple environments have potential for broad deployment in crop breeding to improve rhizosphere community composition, even if the specific taxa associated with those loci are location or location-year specific (Wallace et al. 2018; Wang et al. 2022). The co-localizations of microbial traits between field trials as well as with QTL identified in previous barley rhizosphere microbiome mapping (Escudero-Martinez et al. 2022) lend confidence to the associations reported here.

We particularly detected barley genetic associations with microorganisms that have reported nutrient cycling functions. Nitrospirota bacteria were the most widely represented in the GWAS results (**Fig. 3a**). Members of this taxon carry out nitrification, converting ammonia to a more readily usable and less toxic form of nitrogen for plants (Koch et al. 2015; Vijayan et al. 2021). Thus, these microbes could potentially benefit crops in both under– and over-fertilized conditions, both of which were likely encountered across our field trials (**Table S1**). The loci associated with this microbe (and others) varied between location-years and could be dependent on environmental contexts that include management conditions. Microorganisms involved in nutrient cycling are critical to crop performance (Bargaz et al. 2018). Describing the effects of the quantitative loci underpinning Nitrospirota abundance (**Fig. 3b-h**) could be a step towards selecting for the recruitment of this beneficial bacterium.

Through extensive documentation of plant performance alongside the rhizosphere microbiome characterizations in multiple environments, we linked agronomic and microbial traits to the same genetic loci (**Fig. 4**), expanding beyond previous studies (Escudero-Martinez et al. 2022; He et al. 2024; Meier et al. 2022; Wang et al. 2022). Patterns occurring at such loci suggest that the impact of a genotype on plant fitness is changed upon encountering the right combination of environment and microbial taxa (**Fig. 4a**). A rhizosphere microbial community characteristic occurring during early host development might prime the plant for later success, or the genetic basis for a plant phenotype that impacts the relative abundance of a certain microbial taxon might in turn improve plant fitness. This pervasive co-localization of loci associated with microbial traits and with agronomic traits suggests a link between rhizospheric microbial community composition and crop performance. Such co-localizations can arise from genetic linkage or pleiotropy.

Disentangling the interconnected influences of plant genetics, microbiomes, and abiotic factors to identify cause and effect is a significant challenge (Zancarini et al. 2021), but when addressed will enable selection for microbial traits through breeding. Our results suggest that the general ability to conditionally recruit beneficial taxa under local soil and weather conditions is a viable pathway toward breeding for microbiomes beneficial to crop production. The loci we identified and the aboveground variation we observed together are consistent with conditional, adaptive recruitment by specific barley genotypes in response to local environments. Selecting plant genotypes for conditional recruitment complements other strategies of engineering rhizospheres through inoculation (Mueller and Linksvayer 2022) or selection for specific plant-microbe relationships (He et al. 2024), as it leverages the dynamic and variable microbial taxa that plant genotypes are expected to experience across their adaptive ranges. As knowledge of GxE controls on plant-microbe interactions develops, this strategy may prove more feasible at scale for developing productive and stress tolerant crop varieties.

## Author Contributions

Conceptualization: P.M.E., M.B.K., J.D.S., J.O.E., J.L., Performing Experiments: J.L.W., E.Z.K., A.H.M, P.M.E. Analysis: J.L.W., E.Z.K., A.H.M, J.D.S., J.O.E., J.L., Writing and Revising: J.L.W., E.Z.K., A.H.M, P.M.E., M.B.K, J.D.S., J.O.E., J.L.

## Key Message

Barley genotypes shaped their associated bacterial and fungal rhizosphere communities in both heritable and environmentally dependent ways, connecting host genetic control of microbial recruitment with crop performance.

## Acknowledgements

The authors thank the Hawaii Agricultural Experiment Stations and the Montana State Agricultural Experiment Stations. This work was supported by the United States Department of Agriculture (USDA) National Institute of Food and Agriculture (NIFA) Agriculture and Food Research Initiative (AFRI) award number 2020-67014-32138. Computational efforts were performed on the Tempest High Performance Computing System, operated and supported by University Information Technology Research Cyberinfrastructure at Montana State University. Mention of trade names or commercial products in this publication is solely for the purpose of providing specific information and does not imply recommendation or endorsement by the U.S. Department of Agriculture. The USDA is an equal opportunity provider and employer.

## Statements and Declarations

### Data availability

The raw data for 16S and ITS rRNA sequencing are available in the NCBI Sequence Read Archive database under BioProject PRJNA909304, https://www.ncbi.nlm.nih.gov/bioproject/PRJNA909304. Code is available at https://github.com/Lachowiec-Lab/Barley-genetic-architecture-conditionally-impacts-recruitment-of-rhizosphere-microorganisms/blob/main/

### Ethics approval

Not applicable

### Consent to participate

Not applicable

### Consent to publish

Not applicable

