## Supplemental Figures for "Barley genetic architecture conditionally impacts recruitment of rhizosphere microorganisms and crop performance"

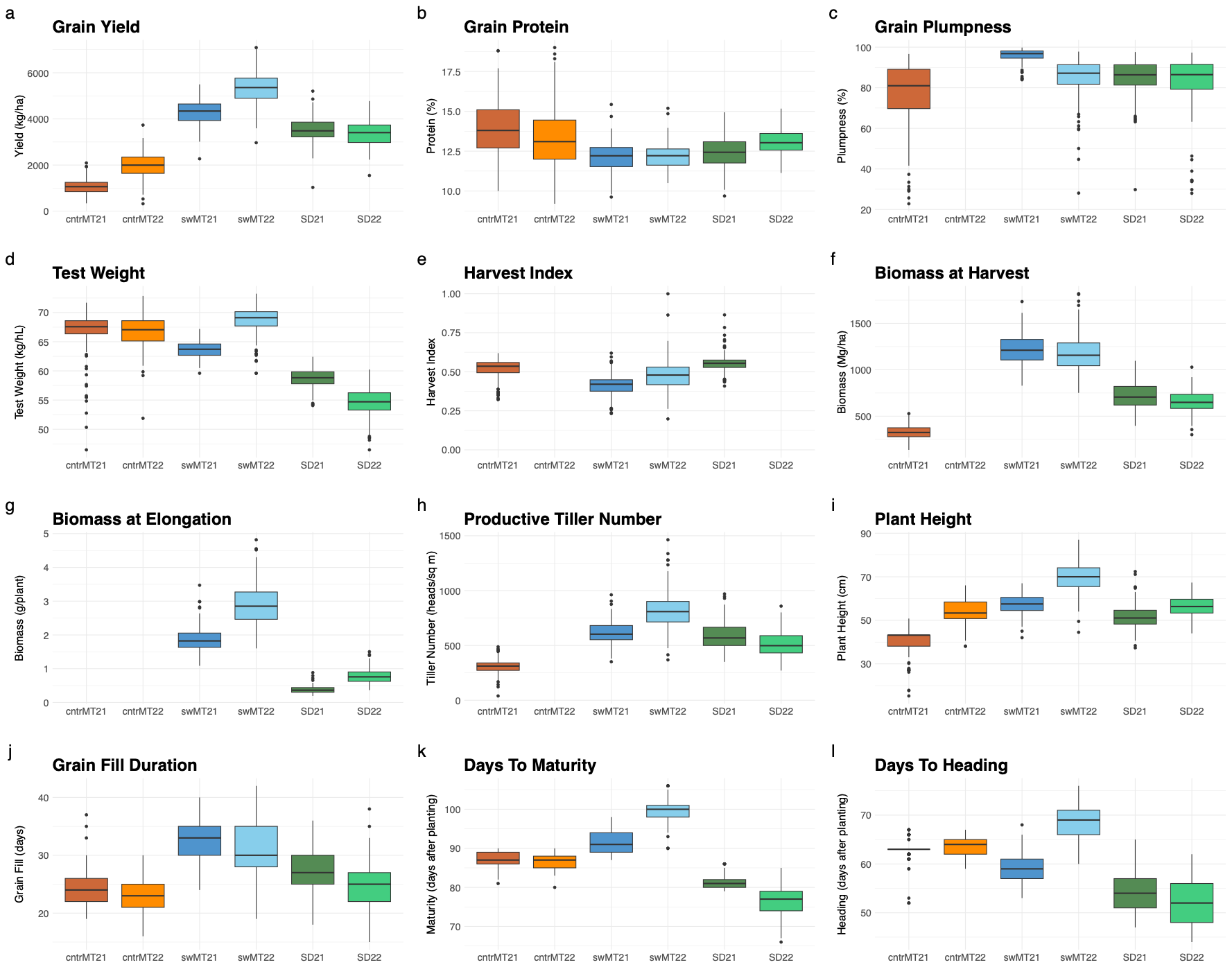

**Figure S1**. Agronomic trait distributions for the S2MET barley population across the six field trials in the adapted range.

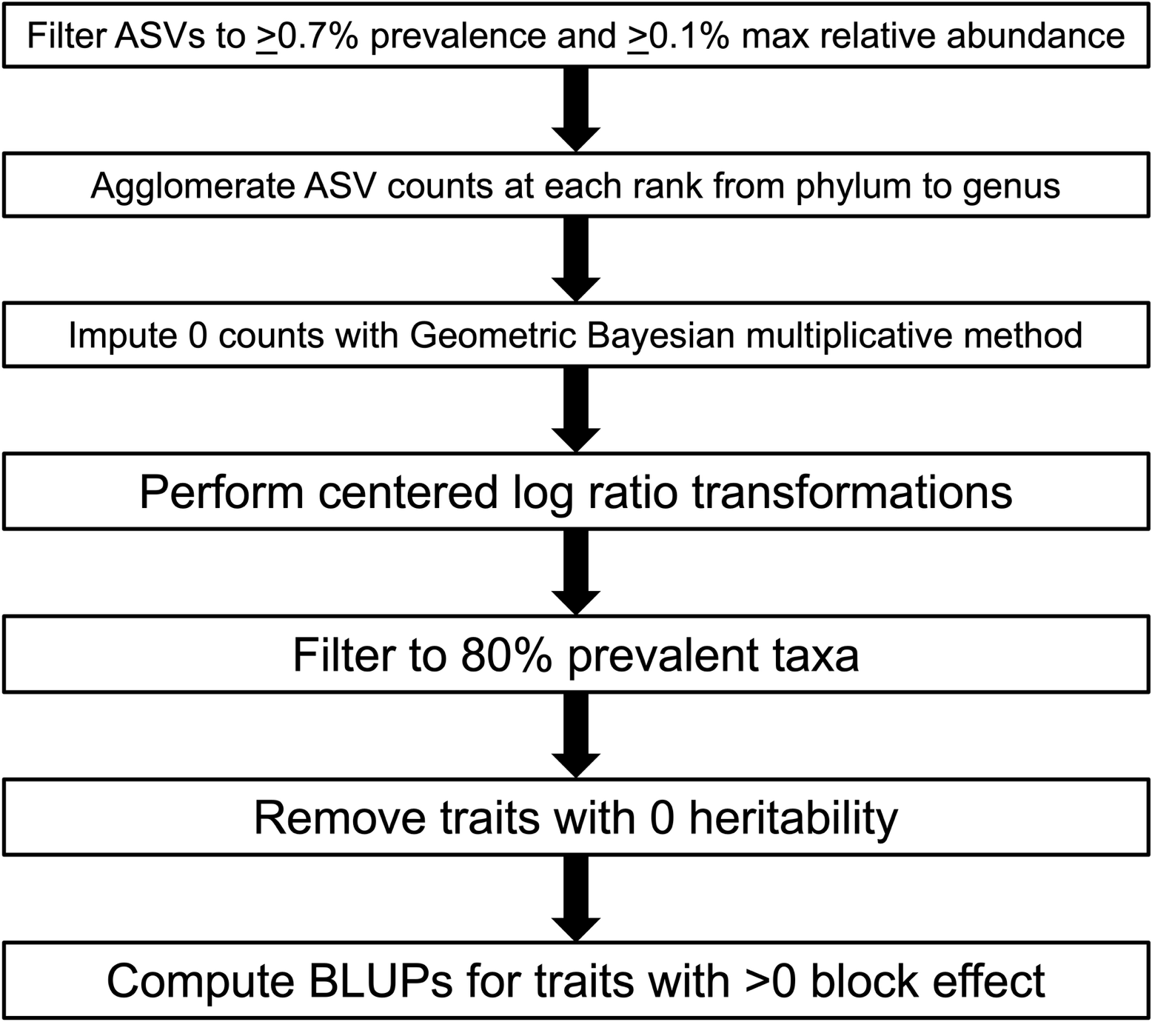

**Figure S2.** Overview of microbial trait preparation for GWAS. These steps were carried out within each location-year dataset separately.
